# Immunization with *Mycobacterium vaccae* NCTC 11659 prevents the development of PTSD-like sleep and behavioral phenotypes after sleep disruption and acute stress in mice

**DOI:** 10.1101/2020.05.07.082859

**Authors:** Samuel J. Bowers, Sophie Lambert, Shannon He, Christopher A. Lowry, Monika Fleshner, Kenneth P. Wright, Fred W. Turek, Martha H. Vitaterna

## Abstract

Because regular sleep disruption can increase vulnerability to stress-related psychiatric disorders, there is a need to explore novel countermeasures to increase stress resilience after inadequate sleep. In this study, we explored the impact of 5 days of intermittent sleep disruption on vulnerability to acute social defeat stress in mice, and investigated the ability of the environmental, immunomodulatory bacterium *Mycobacterium vaccae* NCTC 11659 (MV) to promote stress resilience in that context. We found that mice receiving sleep disruption plus acute stress developed sleep and behavioral phenotypes that had some features of human posttraumatic stress disorder (PTSD) including reduced NREM delta power and increased NREM beta power in post-stress sleep EEG, persistent increases in sleep fragmentation and the REM:Sleep ratio, and behavioral changes. Importantly, immunization with heat-killed MV prevented the development of this phenotype. These results support further research into novel, microbial-based countermeasures to improve health and increase resilience to sleep disruption.

## Introduction

A pattern of repeated nights of inadequate sleep is common in Western, urban societies, and has been shown to lead to a host of physiological problems including metabolic^1^, immunologic^2^, and cognitive^3^ deficits. Disrupted sleep also alters the responsiveness of the hypothalamic-pituitary-adrenal (HPA) axis^4-6^, increases inflammation^7,8^ and potentiates the effects of a chemical stressor in a model of colitis in mice^9^, indicating that the sleep-deprived state may be a stress-vulnerable state. Indeed, there is evidence that short sleep duration is associated with stress-related psychiatric disorders such as depression/anxiety^10^ as well as posttraumatic stress disorder (PTSD) following trauma exposure^11^. Unfortunately, professions with high rates of PTSD such as warfighters, law enforcement, and emergency workers also often find it difficult to get adequate sleep due to the nature of their jobs. There is a need, therefore, to study the relationship between sleep disruption and stress vulnerability and to investigate potential countermeasures to increase resilience to the “double hit” of repeated sleep disruption plus acute stress.

*Mycobacterium vaccae* (strain NCTC 11659) is a non-pathogenic saprophytic bacterium with immunoregulatory and anti-inflammatory properties^12^ and is prevalent in the environment. Bacteria like *M. vaccae* have been labeled “Old Friends” because mammals evolved with frequent exposure to these bacteria, which are thought to play a beneficial role in health (including mental health) by promoting differentiation of regulatory T cells and thereby immunoregulation^13^. Importantly, recent studies have demonstrated that immunization with a heat-killed preparation of *M. vaccae* protects against stressor-induced changes in the gut microbiota^14^, blunts stressor-induced potentiation of chemically-induced colitis^14,15^, prevents stress-induced neuroinflammation and anxiety-like defensive behavioral responses^16,17^, enhances fear extinction when given before or after fear conditioning^18,19^, and prevents a PTSD-like behavioral syndrome in a mouse model of chronic psychosocial stress^14^. Follow-up studies suggested peripheral administration of *M. vaccae* may be exerting its stress-protective effects via promotion of an anti-inflammatory state in the brain and prevention of stress-induced microglial priming^16,20^. It remains unknown, however, what effect *M. vaccae* has on stress-induced changes in sleep, or whether *M. vaccae* immunization can confer stress-protective effects in the context of sleep disruption. The goal of this study was to characterize the impact of repeated sleep disruption, acute stress, and sleep disruption plus acute stress (the double hit) on sleep, physiology, and behavior in mice, and to determine if *M. vaccae* immunization can ameliorate these effects.

## Results

Male C57BL/6N mice (*N* = 115) were randomly assigned to one of eight experimental groups. The experiment followed a 2 x 2 x 2 design (**Figure 1a,b, see Methods**). Mice received subcutaneous injections once per week for three weeks with either a heat-killed preparation of *M. vaccae* (NCTC 11659) or vehicle (borate-buffered saline). After baseline sleep recording, groups were then exposed to a five-day sleep disruption protocol or allowed to sleep *ad libitum*. Immediately after the fifth sleep disruption period, groups were exposed to a 1-hour social defeat or a control manipulation (1-hour novel cage). Behavior in the OLM test was assessed the day after social defeat stress, sleep was recorded throughout the experiment, and tissues were collected at the end of the experiment for analyses (**Figure 1a,b**).

**Figure 1:**
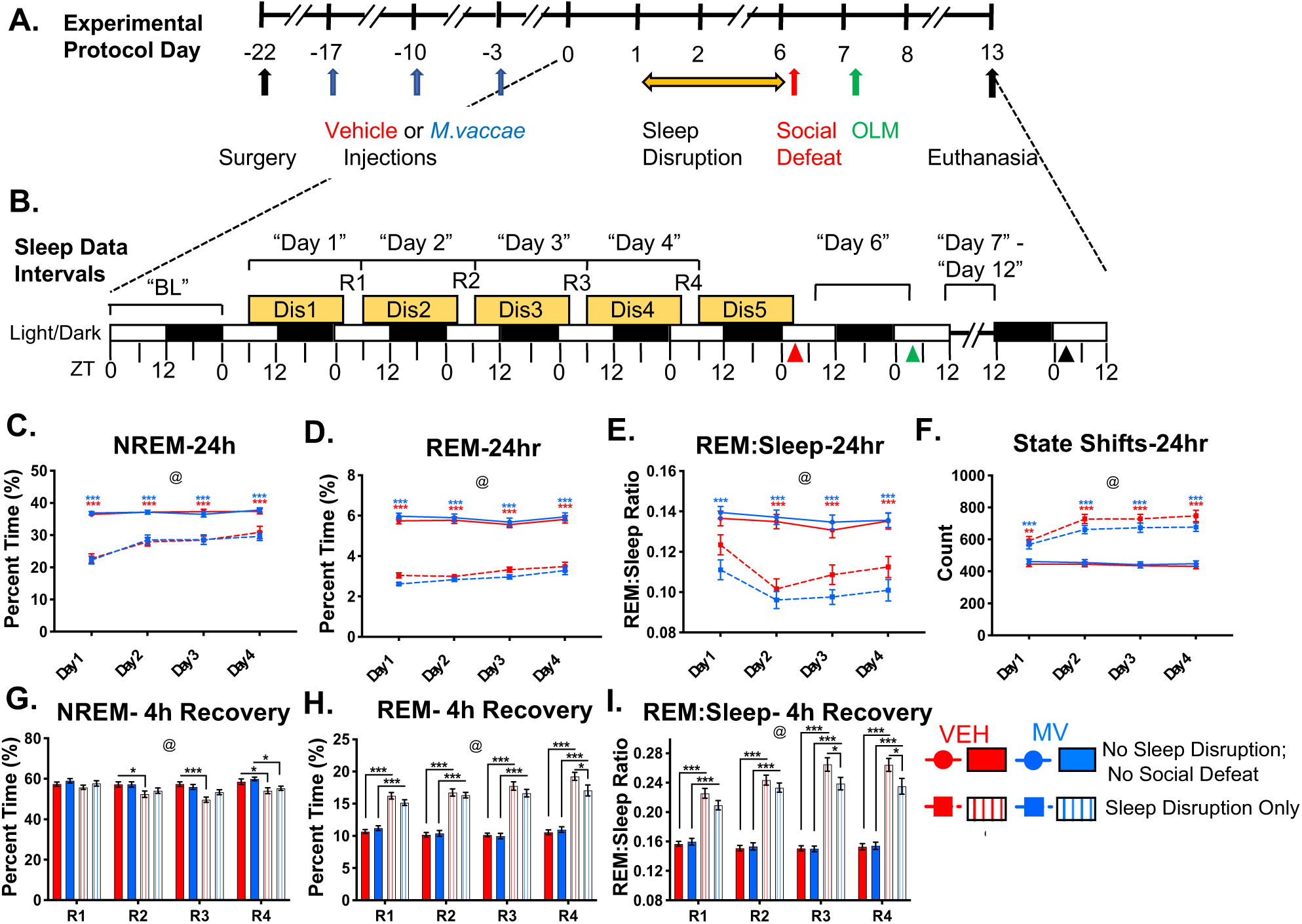
Experimental Timeline and Effect of Sleep Disruption on Sleep in Vehicle-Treated and *M. vaccae*-Treated Mice. (A) Overview of the experimental protocol. Experimental protocol days are depicted along the axis of the timeline, with tick marks indicating ZT0 of that experimental day. (B) Detail of indicated time period that includes the sleep disruption protocol, social defeat stress (red arrow), object location memory (green arrow), and terminal sample collection (black arrow). Tick marks along the bottom indicate zeitgeber time on that day, white/black rectangles indicate 12 h light/dark phases, and yellow rectangles indicated time periods of sleep disruption for the sleep-disrupted groups (ZT6-ZT2). EEG/EMG recording intervals, as used in the forthcoming figures, are indicated above the timeline in quotation marks. (C) NREM, (D) REM, (E) REM:Sleep ratio, and (F) state shifts per 24 h (ZT6-ZT6) during the sleep disruption. (G) NREM, (H) REM, and (I) REM:Sleep ratio during the four hour (ZT2-ZT6) *ad libitum* sleep opportunities. Data are mean + SEM. Symbols: @ *p* < 1.0 x 10-4 (overall effect of “Sleep Disruption”), linear mixed effects model; \**p* < 0.05, \*\**p* < 0.01, \*\*\**p* < 0.001 (Tukey’s post hoc test). Red asterisks indicate vehicle-injected control group vs vehicle-injected sleep-disrupted group at that timepoint. Blue asterisks indicate MV injected control group vs MV injected, sleep-disrupted group at that time point. *n* = 29-30/group. Abbreviations: BL, baseline; Dis, sleep disruption; EEG, electroencephalography; EMG, electromyography; MV, *Mycobacterium vaccae* NCTC 11659 injection; NREM, non-rapid eye movement; OLM, object location memory; R, recovery; REM, rapid eye movement; VEH, vehicle injection; ZT, zeitgeber.

### *M. vaccae* immunization has minimal impact on baseline sleep measured three days after the final injection

Electroencephalogram (EEG) and electromyogram (EMG) recording devices were implanted before the first injection of *M. vaccae* or vehicle (i.e., on day –22), and 24 h baseline sleep was recorded starting three days after the third injection (i.e., on day 0). We observed no group differences between the vehicle- and *M. vaccae*-treated groups in the amount of time spent in non-rapid eye movement (NREM) sleep, rapid eye movement (REM) sleep, the REM:Sleep ratio, or the number of state shifts (**Figure S1a-d**). For *F* statistics and *p* values for all LMM analysis, see **Table S1**.There were no differences between groups in the NREM EEG power spectra or the REM EEG power spectra (**Figure S1e,f**). Overall these results suggest that the three immunizations with *M. vaccae* did not have an appreciable impact on sleep measures under basal conditions.

### The 5-day sleep disruption protocol significantly reduces NREM and REM sleep, and increases sleep fragmentation in both vehicle-injected and *M. vaccae*-injected mice

The sleep disruption protocol used in this study consisted of 5 days of disruption during which a motorized bar at the bottom of the cage disrupted sleep for 20 h/day (ZT6-ZT2), followed by a 4 hour *ad libitum* recovery window (ZT2-ZT6; see **Methods**). EEG/EMG recording during the sleep disruption protocol revealed a significant reduction in 24 h NREM and REM sleep, along with a significant increase in the number of state shifts in sleep-disrupted mice (**Figure 1c-f)**. A close inspection of sleep measures during the 20 h disruption windows showed that the NREM sleep was severely disrupted while the motorized bar was turning. This was characterized by a greater number of bouts of NREM and a reduced median bout duration, along with a reduction of NREM delta (0.5-4 Hz) EEG power and increase in NREM beta (15-30 Hz) EEG power (**Figure S2f-m)**. The mice reached REM sleep extremely rarely during the 20 h sleep disruption window (**Figure S2g**).

Examining the 4 h recovery windows (ZT2-ZT6), sleep-disrupted mice showed a large rebound in the amount of REM sleep during recovery compared to controls (**Figure 1h**). This rebound was similar in magnitude in vehicle-treated and *M. vaccae*-treated groups until recovery window R4, when the percent of time spent in REM was higher in vehicle-treated, sleep-disrupted mice than their *M. vaccae*-treated counterparts (**Figure 1h**). The recovery sleep was REM-dominant, as indicated by a significant increase in the REM:Sleep ratio (**Figure 1i**). Vehicle-injected, sleep-disrupted mice exhibited a greater increase in REM:Sleep than did *M. vaccae*-injected, sleep-disrupted mice during R3 and R4 (**Figure 1i**). This recovery sleep was so REM dominant that it resulted in a reduction in the amount of NREM sleep in the sleep-disrupted groups compared to controls, an effect that increased over time, beginning on R2 in vehicle-treated mice, but not until R4 in *M. vaccae*-treated mice (**Figure 1g)**. NREM sleep during recovery sleep, though reduced in quantity, often featured increased EEG delta (0.5-4 Hz) power (**Figure S2a**). The REM rebound in sleep-disrupted groups during the recovery windows was accompanied by an increase in REM theta2 (8-12Hz) EEG power in vehicle-treated mice at R2-R4 and *M. vaccae*-treated mice at R3-R4 (**Figure S2d**). The sleep disruption protocol also resulted in a plateau in the growth curves of both vehicle-treated and *M. vaccae*-treated mice, with body weights promptly recovering after return to *ad libitum* sleep (**Figure S3**).

### The double hit causes a maladaptive sleep phenotype acutely after social defeat that is prevented by *M. vaccae* immunization

After the fifth 20 h sleep disruption period, half of the mice were exposed to a 1 h social defeat or control manipulation (see **Methods**). Sleep in the immediate aftermath of traumatic events is thought to be important to processing and coping with the negative experience^21-23^. Therefore, upon return to their home cage, EEG/EMG signals were recorded for the next ∼20 hours to examine post-stress sleep (∼ZT7-ZT3, “Day 6”, see **Figure 1a,b**). In vehicle-treated animals, NREM sleep was relatively unchanged in all groups (**Figure 2a**), while the sleep disruption only and double hit groups showed large increases in REM sleep and the REM:Sleep ratio compared to controls (**Figure 2b,c)**. Social defeat increased the number of brief arousals per hour in vehicle-treated animals (overall effect of social defeat, *p* = 0.004), especially in those receiving the double hit (**Figure 2d**). *M. vaccae* immunization altered this phenotype. Among *M. vaccae*-treated mice, there was an increase in NREM after the double hit, relative to mice in control conditions, that was not present in vehicle-treated mice (**Figure 2a**). Furthermore, *M. vaccae* immunization prevented the double-hit induced increase in brief arousals that was seen in vehicle-treated mice (**Figure 2d**).

**Figure 2:**
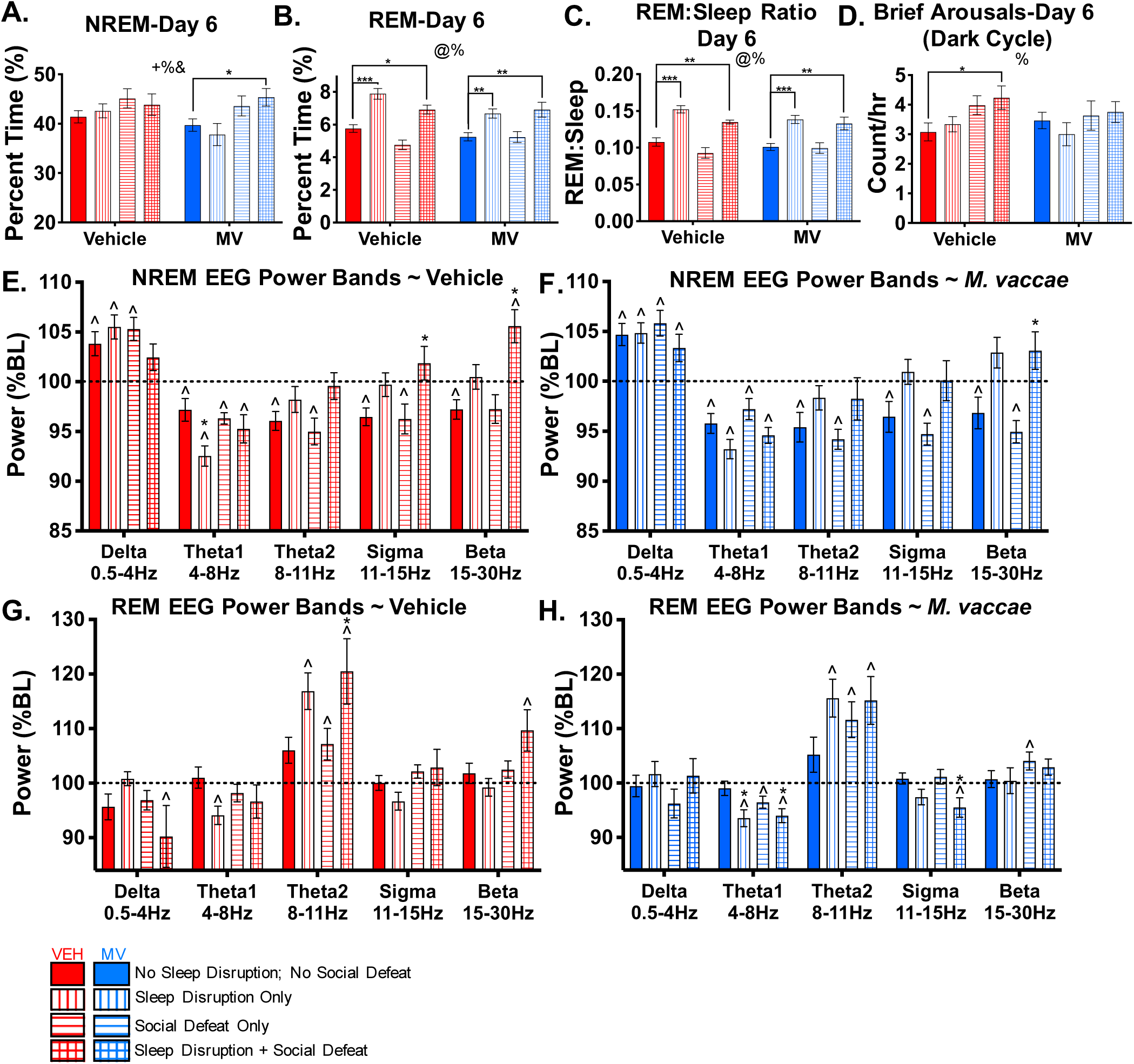
The Double Hit Causes a Maladaptive Sleep Phenotype Acutely After Social Defeat, Which is Prevented by *M. vaccae* Immunization. EEG/EMG sleep recordings were started after social defeat or control manipulation (∼ZT7) and continued until object location memory (OLM) testing began the next morning (ZT3), for a ∼20h window labeled “Day 6”. (A) NREM, (B) REM, (C) REM:Sleep ratio during Day 6. (D) Brief arousals during the dark cycle of Day 6. (E-F) NREM EEG spectral power and (G-H) REM EEG spectral power during Day 6, expressed as a percent of the power during the identical ZT time window during baseline recording. Data are mean + SEM. Symbols in panels (A-D): + *p* < 0.05 (overall effect of “Treatment”); @ *p* < 0.05 (overall effect of “Sleep Disruption”); % *p* < 0.05 (overall effect of “Social Defeat”); and *p* < 0.05 (“Treatment” x “Sleep Disruption” x “Social Defeat” interaction), linear mixed effects model; \**p* < 0.05, \*\**p* < 0.01, \*\*\**p* < 0.001 (Tukey’s post hoc test). Symbols in panels (E-H): ^*p* < 0.05 vs 100% (Wilcoxon Rank-Sum test), \**p* < 0.05 vs Control (Tukey’s post hoc test). *n* = 10-14/group. Abbreviations: BL, baseline; EEG, electroencephalography; EMG, electromyography; OLM, object location memory; MV, *Mycobacterium vaccae* NCTC 11659 injection; NREM, non-rapid eye movement; REM, rapid eye movement; VEH, vehicle injection; ZT, zeitgeber time.

In vehicle-treated mice, the control manipulation, sleep disruption alone, and social defeat alone resulted in an increase in NREM EEG delta power relative to the same time window at baseline (i.e., greater than 100%), accompanied by either a reduction or no change in theta1(4-8 Hz), theta2 (8-12 Hz), sigma (11-15 Hz), or beta (15-30 Hz) power (**Figure 2e**). In vehicle-treated mice receiving the double hit, however, this rebound in NREM delta power, thought to be an adaptive phenomenon^21,22^, was absent (**Figure 2e**). Instead, a significant increase above baseline in NREM beta power was observed (**Figure 2e**). In REM sleep, there was an increase in theta2 EEG power due to sleep disruption, though only the vehicle-injected, double hit group was significantly higher than the control group (**Figure 2g**). Similar to findings in NREM sleep, an increase in REM beta EEG power relative to baseline was only seen in the double hit group (**Figure 2g**).

*M. vaccae* immunization prevented the double hit-induced loss of NREM delta power rebound that was seen in vehicle-injected mice (**Figure 2f**). The increase in NREM and REM beta power relative to baseline that was seen in vehicle-injected, double hit-exposed mice was also not present in *M. vaccae-* treated mice (**Figure 2f,h**). *M. vaccae* groups exhibited largely the same phenotype of NREM theta1, theta2, and sigma power bands as did their vehicle-injected counterparts (**Figure 2f**). Overall, sleep in the immediate aftermath of the double hit (“Day 6”) was characterized by increased brief arousals, a lack of NREM delta power rebound, and increased NREM and REM beta power in vehicle-treated mice, and this phenotype was prevented by *M. vaccae* immunization.

### The double hit causes lasting changes in sleep that are prevented by *M. vaccae* immunization

In order to assess the lasting impact of sleep disruption, social defeat, and the double hit on sleep, *ad libitum* sleep was recorded for six days (day 7-12). Social defeat alone did not cause any lasting changes in sleep measures (**Figure S4**). In vehicle-treated mice, sleep disruption alone and the double hit did not result in a lasting change to NREM sleep compared to control (**Figure 3a**), but REM remained elevated above controls for multiple days (**Figure 3b**). There was not a sustained overall effect of sleep disruption alone on the REM:Sleep ratio in vehicle-treated mice (**Figure 3c**). In the vehicle-treated double hit group, however, the REM:Sleep ratio was elevated compared to control until Day 11 and there was an overall effect of the double hit based on mixed effects linear modeling (**Figure 3c**). This lasting increase in REM:Sleep in the vehicle-injected double hit group was accompanied by a trend (*p* = 0.057) towards a decrease in the average number of epochs of NREM preceding a REM bout (average NREM to REM duration) (**Figure 3d**). Furthermore, the vehicle-injected double hit group displayed a lasting elevation in the number of state shifts per 24 h that was not present in the sleep disruption only group (**Figure 3e**). The NREM EEG beta power was elevated in the double hit group compared to control on Day 7 but returned to control levels thereafter (**Figure 3f**). Importantly, *M. vaccae* immunization prevented all of the aforementioned sleep alterations due to sleep disruption alone or the double hit (**Figure 3**).

**Figure 3:**
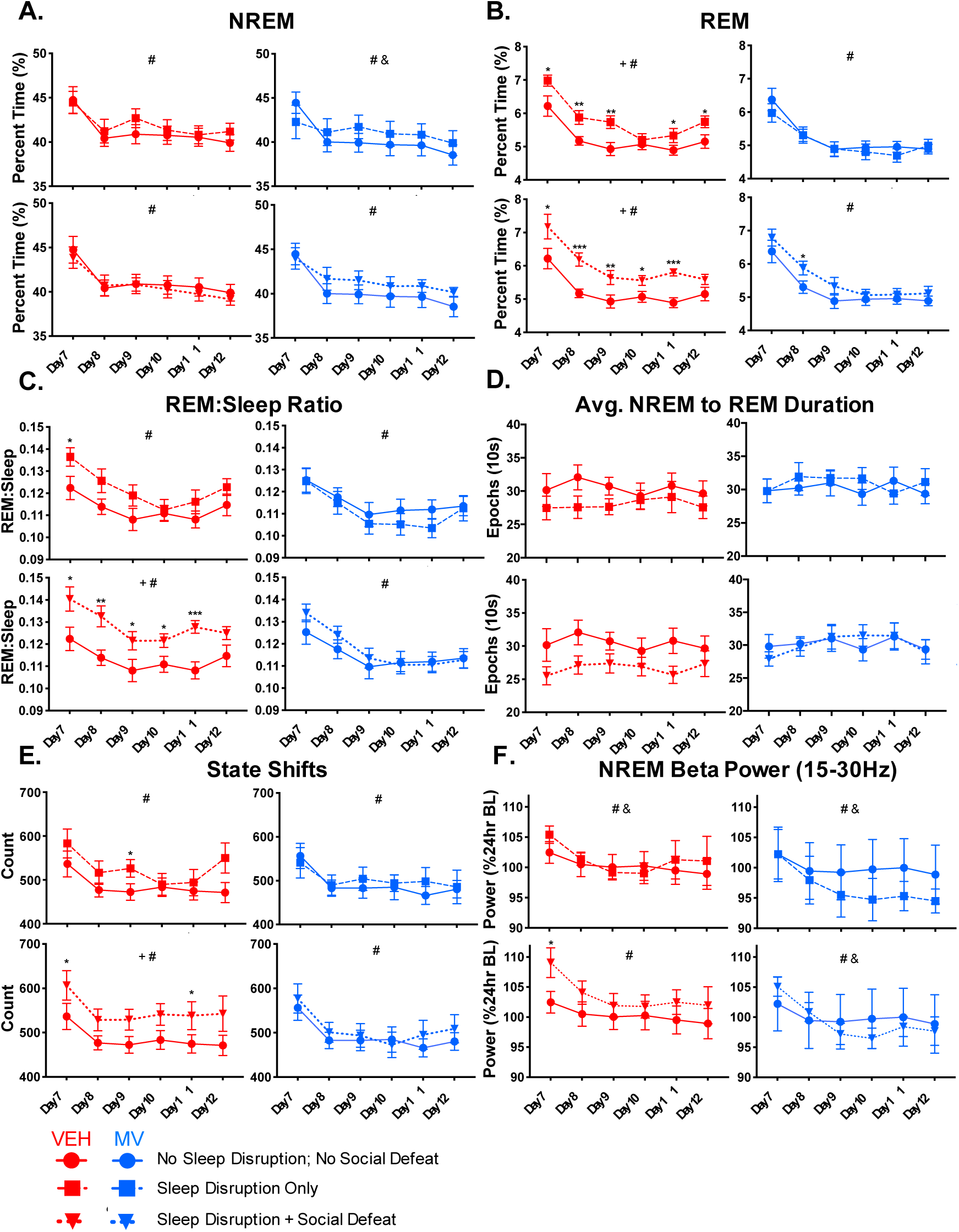
The Double Hit Causes Lasting Changes in Sleep That Are Prevented by *M. vaccae* Immunization. *Ad libitum* recovery sleep was recorded from Day 7 (after object location memory testing) until the end of the experiment (Day 12). 24-h bins of (A) NREM, (B) REM, (C) REM:Sleep ratio, (D) average number of epochs of NREM preceding a REM bout, (E) state shifts, and (F) NREM EEG beta (15-30 Hz) power is reported for control, sleep disruption only, and sleep disruption plus social defeat (double hit) groups. Beta power is reported as a percent of 24-h baseline. Data are mean + SEM. Symbols: + *p* < 0.05 (overall effect of sleep disruption or double hit), # *p* < 0.05 (overall effect of time), & *p* < 0.05 (group x time interaction), linear mixed effects model; \**p* < 0.05, \*\**p* < 0.01, \*\*\**p* < 0.001 (Tukey’s post hoc testing between groups at individual timepoints). *n* = 10-14/group. Abbreviations: BL, baseline; MV, *Mycobacterium vaccae* NCTC 11659 injection; NREM, non-rapid eye movement; REM, rapid eye movement; VEH, vehicle injection.

### The double hit causes hyperlocomotion and anxiety-like behavioral responses that are prevented by *M. vaccae* immunization

The day after social defeat stress, behavior was assessed using the object location memory (OLM) task, which consists of a training session followed by a testing session 90 min later (see **Methods**). Vehicle-treated mice receiving the double hit were hyperlocomotive in the training session compared to controls (**Figure 4a**). Habituation to the testing arena was measured by the percent change in locomotion from one session to the next. While control, sleep disruption only, and social defeat only groups displayed significant reductions in locomotion, mice receiving the double hit did not (**Figure 4c**). *M. vaccae* treatment prevented the double hit-induced hyperlocomotion in the training session (**Figure 4a**), reduced locomotion in the testing session (**Figure 4b**), and prevented the double hit-induced loss of habituation (**Figure 4c**). Anxiety-like defensive behavioral responses during the initial training session were assessed by measuring the percent of time avoiding the center of the arena (see **Methods**). There was an overall effect of social defeat and a sleep disruption x social defeat interaction based on linear mixed effect modeling, as well as a significant reduction in time spent in the center due to the double hit compared to controls in vehicle-injected mice (**Figure 4d**). *M. vaccae*-treated mice showed a reduction in time spent in the center in the social defeat alone group compared to *M. vaccae*-treated controls, but the sleep disruption and double hit groups were unchanged relative to controls (**Figure 4d**). Object location memory is a hippocampal-dependent learning task that is sensitive to stress and anxiety^24^. We found a treatment by sleep disruption interaction, treatment by social defeat interaction, and social defeat by sleep disruption interaction in object location memory using linear mixed effects modeling (**Figure 4e**). While no vehicle-treated groups had location indices significantly different from 50%, *M. vaccae*-treated mice receiving control manipulations, social defeat alone, or the double hit learned the task (**Figure 4e**). *M. vaccae*-treated mice receiving sleep disruption alone showed a *decrease* in location index below 50%, that trended towards statistical significance (one sample Wilcoxon Rank-Sum test vs 50% *p* = 0.08; **Figure 4e**). In summary, the double hit caused a behavioral phenotype consisting of hyperlocomotion and increased anxiety-like behavior in vehicle-treated mice, and *M. vaccae* immunization largely prevented this phenotype and enhanced learning in the OLM task.

**Figure 4:**
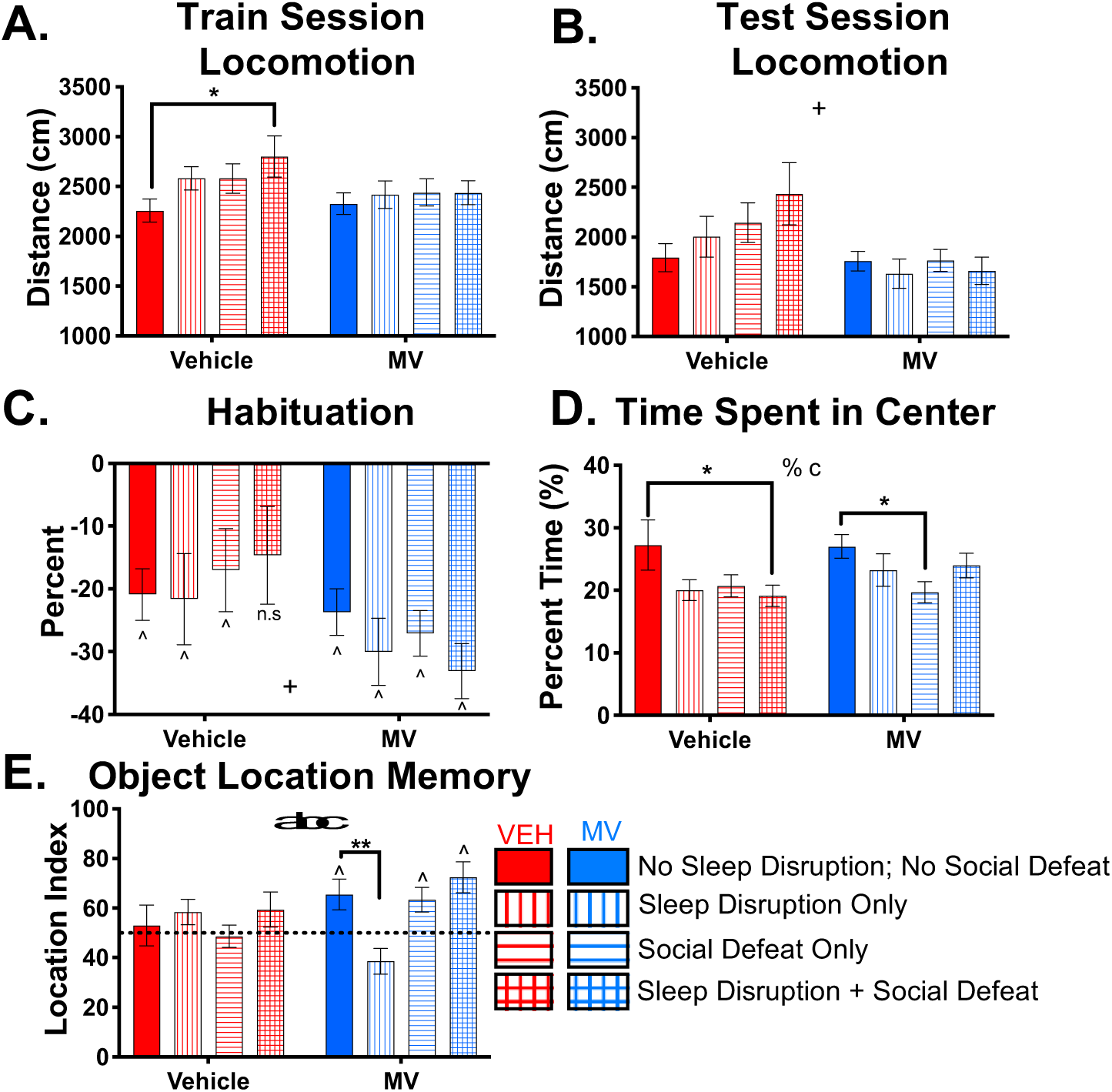
The Double Hit Causes a Behavioral Phenotype That is Prevented by *M. vaccae* Immunization. Approximately 24 h after acute social defeat (or control manipulation consisting of being moved to a clean cage in a quiet room for 1 h), mice were exposed to the object location memory (OLM) task, which consisted of a five-minute training session and a five-minute testing session 90 min later. (A) Total locomotion during the training session and (B) total locomotion during the testing session. (C) Habituation to the testing arena over the two sessions as measured by the percent change in locomotion from the training to the testing session. (D) Percent of the five-minute training session spent in the center of the arena. (E) Object location memory, measured as percent of exploration time devoted to exploring the displaced object during the testing session (Location Index). Significant deviations from 50% indicate learning. Data are mean + SEM. Symbols: + *p* < 0.05 (overall effect of “Treatment”); % *p* < 0.05 (overall effect of “Social Defeat”); a *p* < 0.05 (“Treatment” x “Sleep Disruption” interaction); b *p* < 0.05 (“Treatment” x “Social Defeat” interaction); c *p* < 0.05 (“Social Defeat” x “Sleep Disruption” interaction), linear mixed effects model; \**p* < 0.05, ** *p* < 0.01 (Dunnett’s post hoc test). *n* = 13-15/group. Abbreviations: BL, baseline; MV, *Mycobacterium vaccae* NCTC 11659 injection; NREM, non-rapid eye movement; OLM, object location memory; REM, rapid eye movement; VEH, vehicle injection.

### Sleep disruption, social defeat, and the double hit impacted multiple inflammatory markers. *M. vaccae* treatment ameliorated some, but not all, of these effects

Serum, mesenteric lymph nodes (MLN), and spleen were collected at the end of the experiment, after the 7 days of recovery sleep, to investigate the inflammatory impact of the sleep disruption, social defeat, and the double hit in both vehicle-treated and *M. vaccae*-treated mice (see **Figure 1a,b**). There were no overall effects of treatment, sleep disruption, or social defeat on serum IL-6 or serum monocyte chemoattractant protein (MCP-1; i.e., CCL2) (**Figure 5a,b**). There was an overall effect of social defeat on serum cytokine-induced neutrophil chemoattractant-1 (CINC-1 i.e., CXCL-1) based on linear mixed effect modeling, and post hoc testing revealed a significant increase in CINC-1 due to the double hit compared to controls in vehicle-treated mice (**Figure 5c**). Serum CINC-1 was not increased in the double hit group compared to controls in *M. vaccae*-treated mice (**Figure 5c**). Although linear mixed effect modeling did not reveal an overall effect of treatment, *a priori* comparison of vehicle-treated and *M. vaccae*-treated control groups revealed an increase of serum CINC-1 in the *M. vaccae* group (**Figure 5c**). There was an overall effect of sleep disruption and an overall effect of social defeat on spleen weight based on linear mixed effects modeling, and post hoc testing indicated that the double hit resulted in an increase in spleen weight compared to controls in the *M. vaccae*-treated mice (**Figure 5d**).

**Figure 5:**
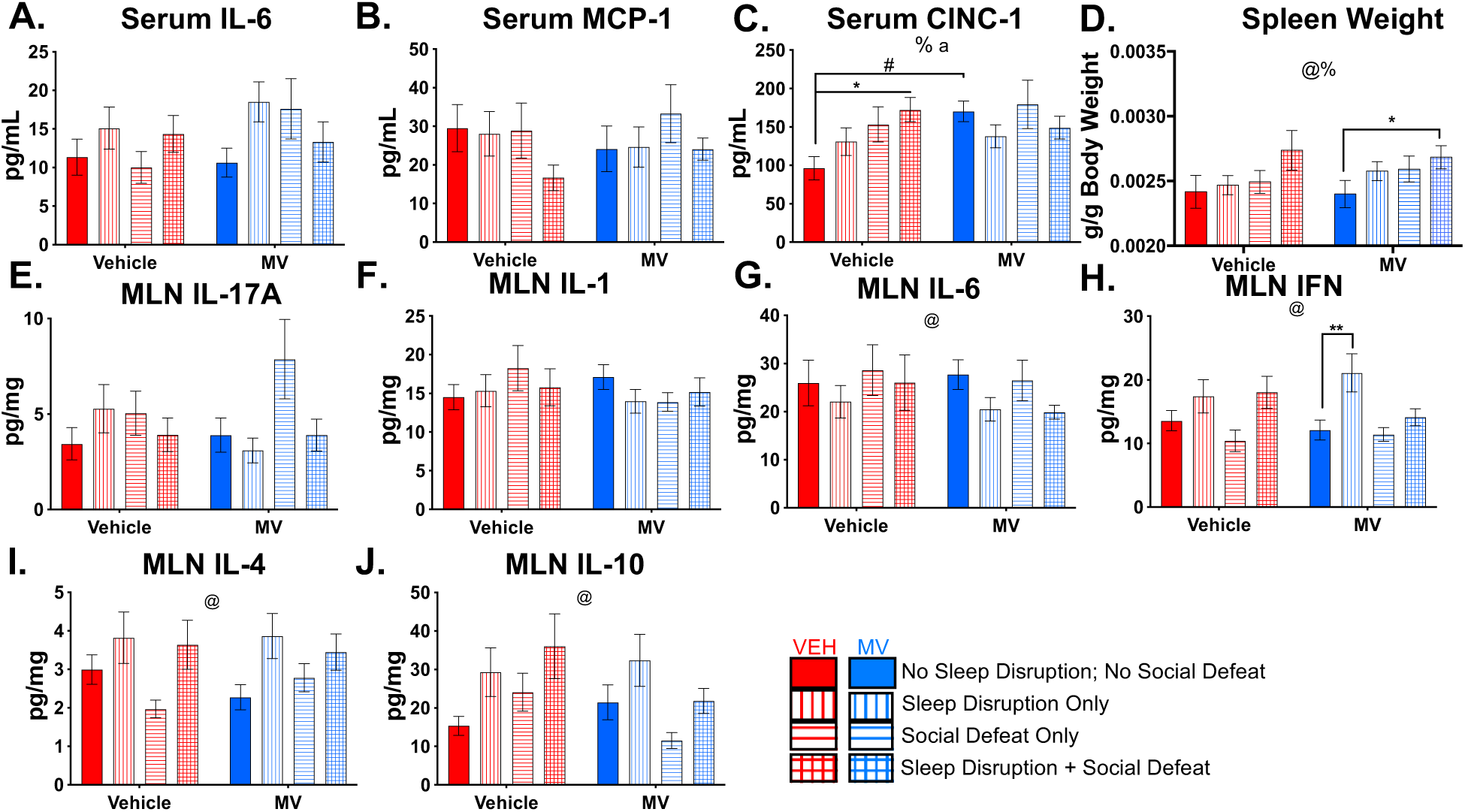
Impact of the Double Hit and *M. vaccae* on Serum Cytokines, Mesenteric Lymph Node Cytokines, and Spleen Weight. At the end of the experiment, serum and other tissues were collected. Serum levels of (A) IL-6, (B) MCP-1 (i.e., CCL2), and (C) CINC-1 (i.e., CXCL-1) were assessed using a magnetic bead multiplex. (D) Spleens were weighed and normalized to body weight. Mesenteric lymph nodes were homogenized and concentrations of (E) IL-17A, (F) IL-1beta (G) IL-6, (H) IFNγ, (I) IL-4, and (J) IL-10 were assessed using a magnetic bead multiplex and normalized to total protein concentration. Data are mean + SEM. Symbols: @ *p* < 0.05 (overall effect of “Sleep Disruption”); % *p* < 0.05 (overall effect of “Social Defeat”); a *p* < 0.05 (“Treatment” x “Sleep Disruption” interaction), linear mixed effects model; \**p* < 0.05, ** *p* < 0.01 (Dunnett’s post hoc test); # *p* < 0.01 (*a priori* Wilcoxon Rank-Sum test). *n* = 11-15/group. Abbreviations: IL, interleukin; CINC-1, cytokine-induced neutrophil chemoattractant-1; MCP-1, monocyte chemoattractant protein-1; MLN, mesenteric lymph node; MV, *Mycobacterium vaccae* NCTC 11659 injection.

The sleep disruption protocol had an impact on multiple cytokines measured in the MLN’s. While there were no overall effects of treatment, sleep disruption, or social defeat on MLN IL-17A or IL-1B (**Figure 5e,f**), there was an overall effect of sleep disruption based on linear mixed effect modeling on MLN IL-6, IFNγ, IL-4, and IL-10 (**Figure 5g-j**). Within these data, the only post hoc comparison to reach statistical significance was an increase in MLN IFNγ seen in the *M. vaccae*-treated sleep-disrupted group compared to *M. vaccae*-treated controls (**Figure 5h**).

### Multiple characteristics of post-double hit sleep predict later changes in sleep, physiology, and behaviour

We then performed pairwise Spearman’s ranked correlation testing of 32 selected variables to examine the relationships among post-social defeat (Day 6) sleep changes, lasting sleep changes (Day 11), and behavioral/physiological changes induced by the double hit (**Figure 6, Figure S5, Table S2**). In vehicle-injected mice, we found 54 associations that were significant after correcting for multiple comparisons (**Figure 6a**). Interestingly, multiple sleep features immediately after social defeat stress (Day 6) correlated with behavioral, sleep, or physiological outcomes days later in vehicle-treated mice. NREM EEG beta power on Day 6 was positively correlated with the REM:Sleep ratio on Day 11 (**Figure 6b**). Other NREM EEG power bands on Day 6 also correlated with outcome measures later in the experiment. NREM delta power on Day 6 correlated negatively with NREM beta power on Day 11 (**Figure S5a**), while NREM theta2 and sigma power both correlated positively with NREM beta power on Day 11 (**Figure S5b,c**). NREM theta2 on Day 6 also correlated negatively with time spent in the center of the arena during the OLM task (**Figure S5d**) and positively with spleen weight at the end of the experiment (**Figure S5e**). Brief arousals on Day 6 were positively correlated with spleen weight at the end of the experiment (**Figure 6c**). Additionally, the REM:Sleep ratio on Day 6 correlated negatively with time spent in the center of the arena during the OLM task (**Figure 6d**) and correlated positively with spleen weight at the end of the experiment (**Figure S5f**).

**Figure 6:**
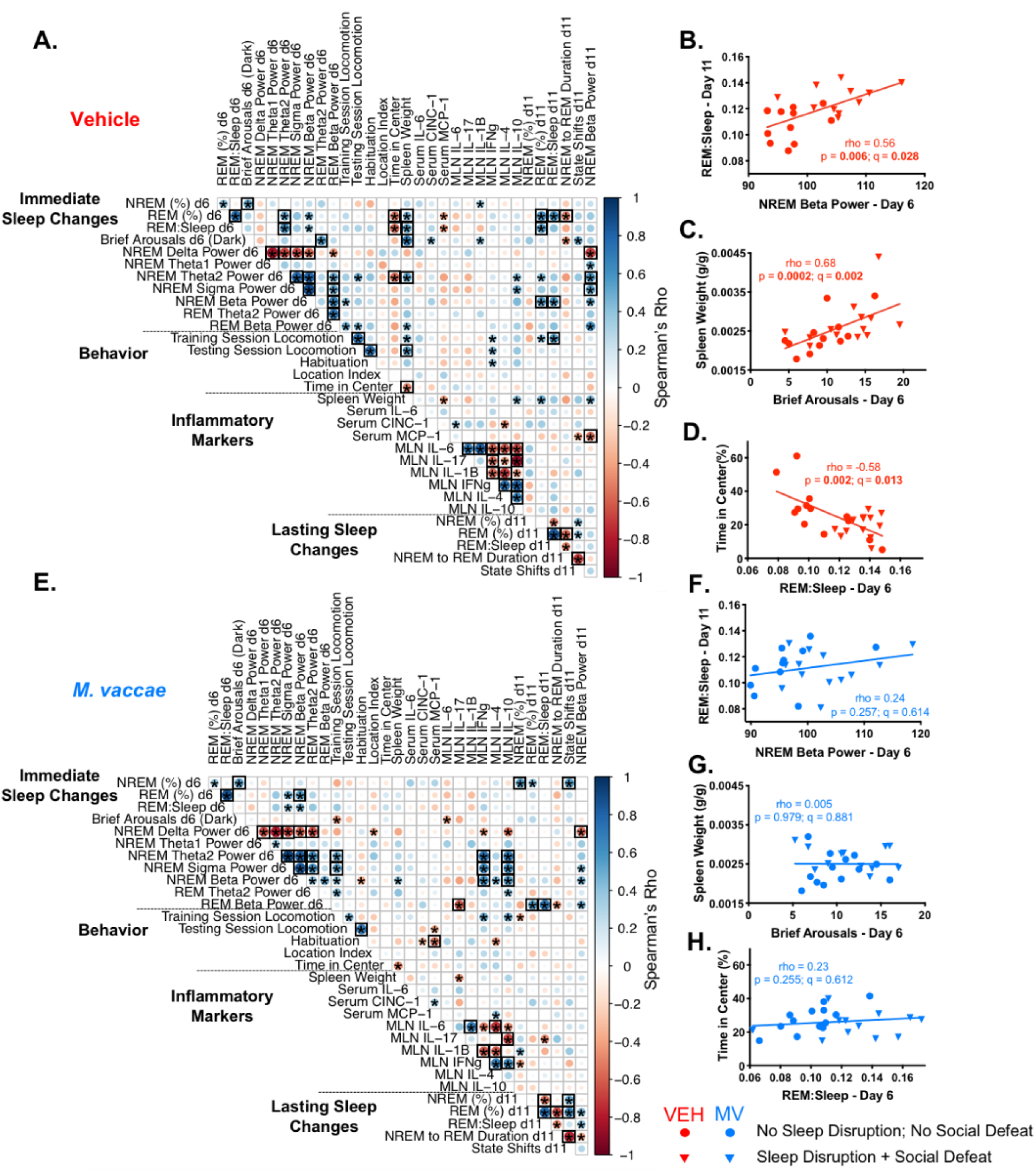
Associations Among Immediate Sleep Changes, Lasting Sleep Changes, Behavior, and Physiological Measures in Vehicle-Treated and *M. vaccae*-Treated Mice. Pairwise Spearman’s rank order correlations of 32 measures of interest from throughout the experiment were computed for (A) vehicle-treated and (E) *M. vaccae* (MV)-treated mice that received either control manipulations or sleep disruption plus social defeat (double hit). The value of the Spearman’s *rho* for each comparison is represented by the size and shade of circle in each square of the correlation plot, with darker blue meaning more positive *rho* and darker red meaning more negative *rho*. Asterisks indicate uncorrected *p* < 0.05, while black boxes indicate *q* < 0.05. REM:Sleep ratio on Day 11 vs NREM EEG beta (15-30 Hz) power on Day 6, spleen weight per gram of body weight at the end of the experiment vs brief arousals during the dark phase of Day 6, and time spent in the center of the arena during the OLM training session vs REM:Sleep on Day 6 are presented in (B-D) vehicle-treated and (F-H) MV-treated mice. *N* = 9-12/group. Abbreviations: EEG, electroencephalography; IFNg, interferon gamma; IL, interleukin; CINC-1, cytokine-induced neutrophil chemoattractant-1; MCP-1, monocyte chemoattractant protein-1; MLN, mesenteric lymph node; MV, *Mycobacterium vaccae* NCTC 11659 injection; NREM, non-rapid eye movement; OLM, object location memory; REM, rapid eye movement; VEH, vehicle injection.

In *M. vaccae*-treated mice, there were 42 significant (*q* < 0.05) correlations between the 32 variables (**Figure 6e**), but the average strength of each tested correlation was significantly weaker than in vehicle-injected groups, as the average Spearman’s rho was significantly smaller (**Figure S5g**). Unlike in vehicle-treated mice, NREM EEG beta power was not significantly associated with the REM:Sleep ratio on Day 11, brief arousals on Day 6 were not associated with spleen weight, and the REM:Sleep ratio on Day 6 was not correlated with behavior the next day (**Figure 6f-h**). In fact, of the aforementioned associations between sleep on Day 6 and subsequent behavior or sleep in vehicle-treated mice, only the negative correlation between NREM delta power on Day 6 and NREM beta power on Day 11 was maintained in *M. vaccae*-treated mice (**Figure S5h**). Instead, there were many associations between EEG power on Day 6 and MLN cytokines. NREM theta2, sigma, and beta power were all positively correlated with both MLN IFNγ and MLN IL-10 (**Figure S5i-n**). NREM theta2 and NREM sigma power on Day 6 were also positively associated with locomotion during the OLM training session in *M. vaccae*-treated mice (**Figure S5o,p**). Finally, while REM beta power was not significantly correlated with any of the other 31 measures after correction for multiple comparisons in vehicle-treated mice, it was positively correlated with the REM:Sleep ratio at Day 11 (**Figure S5q**), and negatively correlated with MLN IL-17 (**Figure S5r**) in *M. vaccae*-treated mice. In summary, multiple features of post-social defeat sleep (including increased NREM beta power, fragmentation, and the REM:Sleep ratio) predicted double hit-induced changes in sleep, behavior, and physiology in vehicle-injected mice. In *M. vaccae*-treated mice, many of these associations were not present, and instead a set of associations between Day 6 sleep EEG power and MLN cytokines emerged.

## Discussion

In this we study tested the hypothesis that sleep disruption induces vulnerability to lasting changes in inflammatory state, sleep, and behavior brought after a “second hit” of acute social defeat stress in mice (the double hit). Furthermore, we investigated immunization with heat-killed *Mycobacterium vaccae* NCTC 11659, a nonpathogenic, soil-derived bacterium with immunoregulatory and anti-inflammatory properties, as a countermeasure to improve resilience to sleep disruption, acute social defeat stress, and the double hit. We found that the double hit resulted in sleep and behavioral phenotypes that were longer lasting and more severe than was seen after sleep disruption or acute social defeat alone. In particular, a maladaptive sleep phenotype immediately after social defeat stress was present only in the vehicle-injected double hit group, multiple features of which predicted changes in behavior, physiology, and sleep disturbances nearly one week later. *M. vaccae* immunization prevented nearly all of the components of the double hit phenotype. Together, these results suggest sleep disruption is a factor promoting vulnerability to lasting effects of acute stress, that inappropriate high frequency EEG power during post-stress sleep is a marker predicting future stress-related sleep impairments, and that immunization with *M. vaccae* is a promising avenue to effectively prevent the development of many of the aspects of the double hit phenotype.

In vehicle-treated mice, five days of repeated sleep disruption potentiated the response to subsequent acute social defeat, resulting in a maladaptive sleep phenotype that was evident within the first 20 hours after social defeat. This phenotype was characterized by increased brief arousals during sleep compared to control mice along with changes to NREM and REM EEG power spectra. Vehicle-treated mice that were mildly stressed by the control manipulation (unplugging from EEG/EMG recording devices and moved in a new cage to a quiet room), those that experienced the sleep disruption protocol alone, and those that experienced 1 h of social defeat alone all displayed rebounds in NREM delta power on Day 6, but the vehicle-treated double hit group did not. NREM delta power is an accepted marker for the homeostatic drive for sleep^25^, and is thought to be important for coping with stressful events^21,22,26,27^. Thus, we interpret an absence of a rebound in delta power after the double hit to be part of a maladaptive change in sleep EEG. The vehicle-treated mice receiving the double hit instead displayed increases in NREM and REM beta power. An increase in high frequency oscillations during sleep is thought to be an indication of cortical hyperarousal, and has been associated with insomnia^28-30^, suicidal ideation^31^, and PTSD^32-34^. Furthermore, two recent studies examining sleep architecture and EEG activity in individuals with PTSD reported the combination of reduced NREM delta power along with increased frontal high frequency power in both NREM^35,36^ and REM^35^ sleep, similar to the pattern seen in this study.

Importantly, along with the maladaptive acute post-stress sleep phenotype, sleep during the 7 days of *ad libitum* recovery in vehicle-injected, double hit mice was altered as well. While increases in NREM and REM sleep in the immediate aftermath of a stressful event are thought to be adaptive^23,37-39^, alterations in sleep that persist long after a traumatic event are thought to be pathological and are commonly observed in trauma- and stressor-related disorders in humans^40,41^. Vehicle-injected mice exposed to the double hit exhibited an increase in sleep fragmentation (as illustrated by an increase in the total number of state shifts), REM sleep, and the REM:Sleep ratio many days after return to *ad libitum* sleep, and these sleep disturbances have been seen in models of highly stress-reactive mice^42^ and in studies of humans with a diagnosis of PTSD. While the exact sleep disturbances seen in PTSD vary^43-46^, a meta-analysis of polysomnographic studies conducted with military veterans and civilians with PTSD found modest changes in multiple sleep parameters including an increase in REM density long after the traumatic event compared to individuals without PTSD^47^. In the present study, the REM:Sleep ratio and measures of sleep fragmentation were increased up to six days into recovery sleep after the double hit in vehicle-injected mice, long after they had recovered to control levels in the sleep disruption alone group. A recent study in humans did not find signs of increased responsivity to acute psychosocial challenge after a single night of sleep deprivation^48^, so it is possible only more protracted or severe disruption protocols like the one in this study increase stress vulnerability. The mechanism by which this repeated sleep disruption increases stress vulnerability warrants further study, and may involve systemic changes that promote a proinflammatory state^7,8^, including changes to the fecal microbiome and metabolome^49^.

We identified multiple features of post-social defeat sleep in vehicle-injected mice that predicted double hit-induced changes to behavior, physiology, and sleep 1-7 days later. These included increased NREM beta power, decreased NREM delta power, increased brief arousals, and increased REM:Sleep ratio, which predicted increases in the REM:Sleep ratio on Day 11, increased NREM beta power on Day 11, increased spleen weight at the end of the experiment, and increased anxiety-like defensive behavioral responses during OLM testing on Day 7. These results suggest features of the first sleep period in the immediate aftermath of a traumatic event (such as signs of cortical hyperarousal paired with lack of delta power rebound) may have utility as biomarkers that can be used to identify individuals with an increased risk of developing lasting trauma-related sleep pathology. However, longitudinal studies in rodent models of PTSD and in humans are required to further evaluate the predictive power of markers during post-trauma sleep and to potentially elucidate the mechanisms involved.

Importantly, *M. vaccae* immunization completely prevented the development of the double hit sleep phenotype in this study (see **Table 1**). The earliest signs of protection in *M. vaccae*-treated mice started during the five-day sleep disruption protocol. These early effects may identify sleep changes conferring stress vulnerability. In vehicle-treated mice, although the amount of REM sleep lost during the 20-hour sleep disruption windows did not change substantially from one day to the next, the subsequent REM rebounds seen during the four-hour recovery windows increased in magnitude across the days of the protocol. This increased REM:Sleep ratio persisted for two days after the sleep disruption protocol in mice who did not receive the second hit of social defeat. A recent study in mice demonstrated that chronic low level stress gradually led to an increase in the REM drive, manifested by an increase in the REM:Sleep ratio, that correlated with measures of HPA axis function^50^. Therefore, the gradual increase in REM during recovery sleep seen in vehicle-injected mice may be indicative of a cumulative stress response process driving REM sleep on top of the homeostatic drive to recover the REM lost in the previous 20 hours. Importantly, this cumulative increase in REM sleep did not occur in *M. vaccae*-treated, sleep-disrupted mice. So, even though the sleep disruption was equally effective in reducing sleep totals in *M. vaccae*-treated mice, their sleep appeared to recover more quickly than in their sleep-disrupted, vehicle-treated counterparts, without any cumulative effects of the five-day protocol. Therefore, *M. vaccae* immunization may modulate the response to the sleep disruption itself, thereby preventing both double hit-induced cortical hyperarousal/sleep fragmentation as well as subsequent protracted increases in sleep fragmentation and the REM:Sleep ratio.

**Table 1:** Similarities between the double hit phenotype and features of human disorders.

| Double Hit Phenotype<sup>1</sup> (Figures 2-5) | Analogous Features of Stress-Related Disorders in Humans | Impact of *M. vaccae* |
| --- | --- | --- |
| Increased brief arousals and state shifts (Figure 2d, 3e) | ► Sleep continuity problems have been observed in many psychiatric disorders<sup>60</sup>, including PTSD<sup>43,60</sup>, depression<sup>60</sup>, and anxiety disorder<sup>60</sup> | **Prevented** |
| Lack of post-stress NREM delta power rebound (Figure 2e) | ► Reduced NREM delta power has been measured in individuals with PTSD<sup>35,36</sup> | **Prevented** |
| Increased NREM beta power (Figure 2d, 3e) | ► Increased sleep EEG beta (15-30 Hz) power has been noted in individuals with insomnia<sup>28-30</sup>, major depression with suicidal ideation<sup>31</sup>, and PTSD<sup>32,33,35,36</sup> | **Prevented** |
| Long term increase in REM (Figure 3b-d) | ► Many<sup>47,61</sup>, but not all<sup>41</sup>, studies in individuals with PTSD show increased REM amounts or REM density<br>► REM sleep elevation has been commonly observed in major depression<sup>60,62</sup> | **Prevented** |
| Hyperlocomotion during OLM, reduced habituation, reduced time in the center of the training arena (Figure 4a-d) | ► Psychomotor agitation and/or hypervigilance is common in individuals with PTSD<sup>63</sup> | **Prevented** |
| Increased serum CINC-1 (Figure 5c) | ► Increased proinflammatory cytokines have been observed in individuals with insomnia<sup>64</sup>, anxiety disorders<sup>65</sup>, depression<sup>66</sup>, and PTSD<sup>65,67-69</sup>, but CINC-1 in particular has not been affiliated with any of these disorders in humans<sup>70</sup>. | **Unclear**<br>CINC-1 was not increased compared to control in MV-double hit group, but there was an overall increase in CINC-1 due to MV. Further, many proinflammatory changes due to sleep disruption were seen in both treatment groups. |

<sup>1</sup>The ‘double hit phenotype’ was defined as the collection of measures that were altered in sleep-disrupted, socially-defeated (double hit) vehicle-injected mice compared to controls (left column). Many features of this phenotype are similar to those seen in various human disorders, particularly those that are often affiliated with stress or trauma, including insomnia, depressive disorders, anxiety disorders, and PTSD. The third column indicates which of these phenotypic features was prevented by immunization with MV. Abbreviations: CINC-1, cytokine-induced neutrophil chemoattractant 1; EEG, electroencephalography; MV, *Mycobacterium vaccae* NCTC 11659; NREM, non-rapid eye movement sleep; OLM, object location memory; PTSD, posttraumatic stress disorder; REM, rapid eye movement sleep.

Another aspect of the double hit phenotype that was prevented by *M. vaccae* was a change in behavior observed during OLM testing. *M. vaccae* prevented the double hit-induced increase in locomotion in both the training and testing sessions. Hyperlocomotion has been observed in models of chronic mild stress^51,52^, and, along with the signs of cortical hyperarousal during sleep, may be indicative of increased central arousal and hypervigilance, both of which are seen in PTSD. *M. vaccae* also enhanced two measures of memory during OLM testing. First, it maintained habituation to the OLM chamber that was lost after the double hit in vehicle-treated mice, again indicating prevention of a persistent state of arousal and hypervigilance. Second, *M. vaccae*-treated mice successfully distinguished between the moved and not-moved objects in the OLM task while no vehicle-injected groups were able to do so. Object location memory is particularly sensitive to anxiety and prior stress^24^, so it is possible that any experimental manipulation, even the control manipulation used in the place of social defeat, was stressful enough to block learning in the vehicle-treated mice. It is unclear from these data alone whether the improvement in OLM memory seen in *M. vaccae*-treated groups was due to resilience to stressful experimental manipulations or due to enhancement of hippocampal memory *per se*. Recent studies have demonstrated that *M. vaccae* enhanced fear extinction in the fear-potentiated startle paradigm in rats whether administered before^19,53^ or after^18^ fear conditioning, and that it prevented memory impairments in a model of post-operative cognitive dysfunction^20^. However, the impact of *M. vaccae* immunization on various forms of memory in a stressor-independent paradigm has not been explicitly studied and deserves follow-up investigation.

The exact mechanisms of *M. vaccae’s* stress protective properties are unknown, but recent work supports the hypothesis that the anti-inflammatory and immunoregulatory properties of the bacterium are key contributors, both centrally and peripherally^14,54^. This study offers some data to support this hypothesis as *M. vaccae* immunization modified the effect of sleep disruption on serum CINC-1 seven days after the sleep disruption ended. However, *M. vaccae* treatment did not attenuate the sleep disruption-induced increases in serum IL-6, MLN IFNγ, MLN IL-4, or double hit-induced increased spleen size seen at the end of the experiment, indicating that some of the inflammatory impact of the double hit was still present. A recent study in mice observed increases in serotonergic activation in the dorsal raphe nucleus acutely after *M. vaccae* injection^55^. Another study in rats observed increases in anti-inflammatory IL-4 expression in the hippocampus eight days after *M. vaccae* administration, along with a reduction in measures of microglial priming^16^. Thus, modulation of serotonergic networks and reduced neuroinflammation may be mechanisms by which *M. vaccae* prevents double hit-induced changes in sleep and behavior, but further studies are required to test this hypothesis directly.

Some aspects of the study may limit interpretation of some results. First, the measures of inflammatory response were taken at the end of the experiment, a full seven days after social defeat. This makes measurement of markers of the inflammatory response to the double hit difficult to interpret. Follow-up studies examining physiological changes directly after the social defeat stress, when cortical hyperarousal was at its peak in vehicle-treated, double hit-exposed mice, could help elucidate the mechanism by which *M. vaccae* prevents the development of the double hit phenotype. Second, the OLM task alone does not fully characterize the behavioral impact of the double hit. Performing multiple behavioral assessments would have prevented the acquisition of undisturbed sleep, one of the most important outcome measures of this study. Evidence of the sleep loss induced by behavioral testing can be seen in this dataset, as the OLM resulted in an increase in sleep the next day in non-sleep disrupted, non-socially defeated control mice on Day 7 compared to the other days of recovery sleep. Third, this study does not address questions about the duration of the double hit phenotype or the duration of *M. vaccae*-induced protection. Follow up experiments that include a time series study will have to be performed before exploring the use of *M. vaccae* in humans.

Taken together, these results indicate that the sleep-disrupted state can be a stress vulnerable state whereby physiological and psychological effects of acute stress may be more severe and longer lasting. In this context, our data support the hypothesis that cortical hyperarousal during post-stress sleep may be an early marker of long-lasting sleep disturbances after a traumatic event, which potentially would be useful criteria for initiation of sleep-targeted interventions before they become severe. Follow-up studies investigating the predictive potential of increased beta power during sleep are warranted. While simply avoiding repeated bouts of sleep disruption would be one approach to improve stress resilience, there are many professions, such as warfighters in combat and emergency responders, which, by the nature of their work, will unavoidably be exposed to intermittent periods of sleep disruption. Therefore, another important conclusion from this study is that immunomodulatory bacteria, and *M. vaccae* NCTC 11659 in particular, have therapeutic potential to improve health in the context of sleep disruption, a common problem in Western urban societies today.

## Materials and Methods

### Animals and experimental design

Five cohorts of 24 seven-week-old male C57BL/6N mice (Charles River Laboratories, USA; *n* = 24 from Charlotte, NC; *n* = 96 from Kingston, NY) were used for this experiment (*N* = 120). A total of 5 animals did not complete the experimental protocol and were thus eliminated from all analyses, for a total *N* = 115. No explicit power analysis was used, the sample size was determined in to ensure adequate number of biological replicates of the main outcome measure: sleep recordings (target of *n =* 10-12). The experiment consisted of 8 experimental groups in a 2 x 2 x 2 design (Vehicle vs *M. vaccae* injection, *ad libitum* sleep vs sleep disruption, no social defeat vs social defeat). Experimental groups were balanced in each cohort. Mice were group housed until EEG/EMG implant surgery, after which they were individually housed until the end of the experiment. After EEG/EMG surgery, mice were assigned to experimental groups randomly, but effort was made to ensure prior cagemates were in different groups. Male CD1 retired breeder mice (Charles River Laboratories, USA) were used as aggressor mice in the social defeat model (see below). All mice were maintained on a 12:12 L:D cycle at room temperature (23 + 2 °C) with food and water available *ad libitum* throughout the experiment. All protocols were approved by the Northwestern Institutional Animal Care and Use Committee. The light source in the sleep recording chambers was two 14 W fluorescent bulbs (soft white, 3000 K), resulting in an average light intensity of ∼500 lux inside the cylindrical sleep disruption cage. Zeitgeber Time (ZT) is defined as the number of hours after the onset of the light period (light onset = ZT0).

### EEG/EMG implantation surgery

One week after arrival, and 5 days prior to the first injection (i.e., on day –22), mice were implanted with electroencephalographic/electromyographic (EEG/EMG) sleep recording devices (Pinnacle Technologies, Lawrence, KS, USA). Surgical procedures were performed using a mouse stereotaxic apparatus with standard aseptic techniques in a ventilated, specially-equipped surgical suite. Anesthesia was induced by intraperitoneal (i.p.) injection of a cocktail of ketamine HCl (98 mg/kg; Vedco Inc, St. Joseph, MO, USA) and xylazine (10 mg/kg; Akorn Inc, Lake Forest, IL, USA) before surgical implantation of a headmount, which consisted of a plastic 6-pin connector connected to four EEG electrodes and two EMG electrodes. Four stainless steel screws serving as 2 EEG leads and grounds were screwed into the skull with one EEG lead located 1 mm anterior to bregma and 2 mm lateral to the central suture, and the other at 1 mm anterior to lambda and 2.5 mm lateral to the central suture. The exposed ends of two stainless steel Teflon-coated wires (0.002 in. in diameter) serving as EMG leads were then inserted into the nuchal muscles using a pair of forceps. The entire headmount was then sealed by dental acrylic and a single skin suture at the front and the back of the implant was placed to close the incision. Heat support was provided until recovery from anesthetic by placing animals on a circulating water blanket. Subcutaneous injection of analgesic meloxicam (2 mg/kg; Norbrook Laboratories, Northern Ireland) was given to the animals at the time of surgery and once more on the following day.

### Mycobacterium vaccae administration

Heat-killed preparations of *M. vaccae* NCTC 11659 were provided by BioElpida (Lyon, France; batch ENG#1). Injections of *M. vaccae* (0.1 mg, 0.1 mL) or vehicle (sterile borate-buffered saline) were administered subcutaneously to the rear-left flank once per week for three weeks. The first injection was given 5 days after EEG/EMG implant surgery, and the third injection was given 3 days before baseline sleep recordings (i.e., days –17, –10, and –3; **Figure 1a**).

### Sleep recording and analysis

One week prior to the sleep disruption protocol (day –6), mice were moved into cylindrical sleep recording cages (25 cm in diameter and 20 cm tall, Pinnacle Technologies, Lawrence, KS, USA) within individual acoustically-isolated and Faraday-shielded chambers and the headmount was connected to the transmission tether. Cages had corncob bedding and food/water available *ad libitum*. Five days were allowed for acclimation to the tether before baseline sleep was recorded (i.e., starting at ZT0 on day 0). The sleep recording windows were divided into the following segments (see **Figure 1a,b**): “**Baseline**” was the 24 h period beginning 30 h prior to the sleep disruption protocol (i.e., starting at ZT0 on day 0). During the sleep disruption protocol, 24 h bins (ZT6-ZT6) were reported as “**Day 1**-**Day 4**”, and contained the 20 h disruption (**Dis:** ZT6-ZT2) followed by the 4 h recovery (ZT2-ZT6) window. This time period was further divided into the 20 h windows during which the motorized bars were on (“**Dis1-Dis5**”) and the 4 h recovery windows (“**R1-R4**”). At the end of the fifth 20 h sleep disruption session, all mice were unplugged from their EEG/EMG tethers for social defeat (or control manipulation, see below). Upon return to home cages, sleep recording resumed for ∼20 h (∼ZT7-ZT3) until mice were unplugged from their EEG/EMG tethers for testing in the OLM test. This segment was labeled “**Day 6**”. After OLM behavior testing was complete, EEG/EMG tethers were again plugged in and sleep was recorded for 6 days until the end of the experiment (“**Day 7-Day 12**”).

Data were collected using Pinnacle Acquisition software (Pinnacle Technologies), then scored as non-rapid eye movement sleep (NREM), rapid eye movement sleep (REM), or wake in 10 second epochs using machine learning-assisted sleep scoring software developed in the Turek/Vitaterna laboratory^56^. The initiation of a bout of NREM, REM, or wake was defined by the occurrence of two consecutive epochs of NREM, REM, or wake (respectively). A bout was terminated when a bout of another state occurred. Sleep bouts were initiated by two consecutive epochs of a sleep state (NREM or REM) and were only terminated when a wake bout occurred. A brief arousal was defined as a single epoch of wake within a sleep bout. The average NREM to REM duration was defined as the average number of epochs of NREM preceding each REM bout. The delta power band was defined as 0.5-4 Hz, theta1 as 4-8 Hz, theta2 as 8-11 Hz, sigma as 11-15 Hz, and beta as 15-30 Hz. Relative power was calculated as the raw power (uV^2) in a particular band divided by the total power in all bands. Power was then reported as a percent of baseline to reduce inter-individual variability.

### Sleep disruption protocol

After 7 days of acclimation to the sleep chambers and baseline sleep recordings, half of the mice entered the sleep disruption protocol. All cages had corncob bedding and food/water available *ad libitum* throughout the protocol. Sleep disruption was achieved using a commercially available system integrated into the chambers (Pinnacle Technology), which simulates the gentle handling technique via a rotating metal bar (22 cm in length) attached to a post at the center of the cage. For the sleep disruption period, the bar’s rotation speed was set at 7 rotations per minute with reversals of rotation direction (i.e., clockwise vs. counterclockwise) set to occur at semi-random intervals of 10 ± 10 seconds. The bar was programmed to rotate for 20 hours per day (ZT6-ZT2), and was stationary from ZT2-ZT6, for 5 days. Experimenters visually inspected mice at regular intervals during the sleep disruption windows to ensure that the bar mechanism was functioning properly and that the sleep-disrupted mice were awake. Control animals were placed in identical cages with bars that remained stationary throughout the experiment.

### Social defeat protocol

Male CD1 retired breeder mice were singly housed in large (44 cm x 24 cm x 21 cm) polycarbonate cages and were screened for aggressive behavior before the experiment. Mice that started to injure their opponents by harmful bites during screening were not used for the social defeat procedure. On the day of the social defeat stressor (between ZT2-ZT4), each experimental mouse was introduced into the cage of an aggressor. As soon as the aggressor mouse attacked and defeated the intruder mouse (upper limit of attack latancy was 30 seconds), the intruder was covered with a 10.2 cm x 12.7 cm x 10.2 cm metal mesh cage while still inside the aggressor cage, and left for 1 h. Control animals not receiving social defeat were also unplugged, and placed in a new cage with clean woodchip bedding for one hour (novel cage as a control manipulation”).

### Object location memory task

The object location memory (OLM) task is a hippocampal-dependent memory task^24^ that is sensitive to stress^57,58^. All groups of mice underwent testing in the OLM task the day after social defeat (or control manipulation), beginning at ZT4. Mice were placed in a dimly lit (∼50 lux) 53 cm x 53 cm x 30 cm arena containing two identical cylindrical black caps on the same side of the arena. Mice were allowed to explore the arena for 5 minutes and after 90 minutes in their home cage were allowed to explore the arena again, with one object moved. Exploration of the moved object more than the non-moved object is considered evidence of successful acquisition of contextual memory, and is denoted by a “location index” (100 x {time exploring moved object/total time exploring either object}) significantly different from 50%. In this experiment, location index was quantified during the first 3 minutes of the testing session. LimeLight (Actimetrics, Wilmette, IL, USA) behavioral software was used to track the animal’s path traveled within the open field over time. De-identified video files were scored by two experimenters blind to treatment or stress manipulation and average location indices were reported. Location index inter-scorer reliability was high (*rho* = 0.96, *p* < 0.0001).

### Tissue collection

Six days after OLM testing (day 13), mice were euthanized via rapid decapitation between ZT4-ZT6 and trunk blood was collected in Z-gel serum tubes (Sarstedt AG & Co, Nümbrecht, Germany). The blood was centrifuged at 7500 rpm for 5 min, and serum was collected and stored at –80 °C until analysis. Spleens were dissected and weighed. Mesenteric lymph nodes were collected and quickly frozen at –80 °C until processing for cytokine analysis (see below).

### Cytokines

Serum cytokines and chemokines were assessed using a customized magnetic bead multiplex (Millipore-Sigma, Burlington, MA, USA) containing: 1) interleukin 6 (IL-6); 2) monocyte chemoattractant protein 1 (MCP-1); and 3) cytokine-induced neutrophil chemoattractant-1 (CINC-1/CXCL-1). Serum samples were diluted 1:2 per the manufacturer’s instructions and assayed in duplicate. Cytokines were also measured in mesenteric lymph node extract. Lymph nodes were homogenized in tissue extraction reagent (ThermoFisher, Waltham, MA, USA) using a glass bead sonicator (Diagenode, Denville, NJ, USA). Tubes were centrifuged and the protein extract was aliquoted for multiplex analysis. Total protein was quantified in the extracts (Pierce BCA, ThermoFisher) and used to normalize results. Extracts were diluted 1:5 and cytokine concentrations were determined using a customized magnetic bead multiplex (Millipore-Sigma) containing: 1) IL-6; 2) IL-10; 3) IL-4; 4) interferon gamma (IFNγ); 5) IL-1β; and 6) IL-17A. Assays were conducted using duplicate samples (technical replicates).

### Statistical analysis and software

All detailed statistical results can be found in **Tables S1-S2**. All graphs depict the mean + SEM. Sleep, behavior, and cytokine data were analyzed using linear mixed effects models using the lmer() and anova() functions in the lme4 package of RStudio (version 1.0.136, RStudio Inc, Boston, MA, USA). Models were tested for effects of treatment, sleep disruption, and social defeat, accounting for random effects of cohort (and plate number for cytokine data). Where appropriate, an effect of time was added to the model, along with terms accounting for repeated measures (see **Table S1**). When an overall effect on LMM was detected, post hoc comparisons were performed using the emmeans() function, with the Tukey or Dunnett correction for multiple comparisons when noted in the text. Two-sided Wilcoxon Rank-Sum testing was performed in RStudio. Spearman’s Rank Based correlation analyses were performed using the rcorr() function within the Hmisc package in RStudio, and visualized using the corrplot package. *P* values from the correlation analyses were adjusted for multiple comparisons using the qvalue package in RStudio (see **Table S2** for rho, *p*, and *q* matrices). All other graphs/figures were generated using GraphPad PRISM (version 7.0a; GraphPad Inc., San Diego, CA, USA).

Outlier testing was performed on all datasets in GraphPad PRISM. For the sleep dataset, when individual observations were detected as extreme outliers using the Grubbs’ test^59^ (two-tailed alpha = 0.05), the EEG/EMG raw data files were visually inspected, and those observations were excluded from analysis if signal noise was deemed to interfere with accurate stage scoring or power analysis. This amounted to 0.62% of the sleep-related datapoints across the entire experiment (117/18,865). Extreme outliers detected in the behavioral datasets with the Grubbs’ test and verified to be due to software tracking errors were excluded (*n* =2/115 mice, 6/460 datapoints). Cytokine and chemokine multiplex datasets contained multiple extreme outliers, and those identified using the ROUT test (*Q* = 1%) were removed from analysis: 1) serum IL-6, *n* = 7; 2) serum CINC-1, *n* = 1; 3) serum MCP-1, *n* = 2; 4) MLN IL-6, *n* = 6; 5) MLN IL-17A, *n* = 8; 6) MLN IL-1β, *n* = 2; 7) MLN IFNγ, *n* = 3; 8) MLN IL-4, *n* = 4; 9) MLN IL-10, *n* = 8.

## Acknowledgements

The authors would like to acknowledge Chris Olker and Eun Joo Song for assistance scoring sleep and Dr. Peng Jiang for guidance throughout the analysis. Serum and MLN assays were enabled by the Comprehensive Metabolic Core at Northwestern University. This work was supported by ONR MURI Grant #N00014-15-1-2809 and NIH Training Grant T32 HL007909.

## Supplementary Figures and Legends

**Figure S1:**
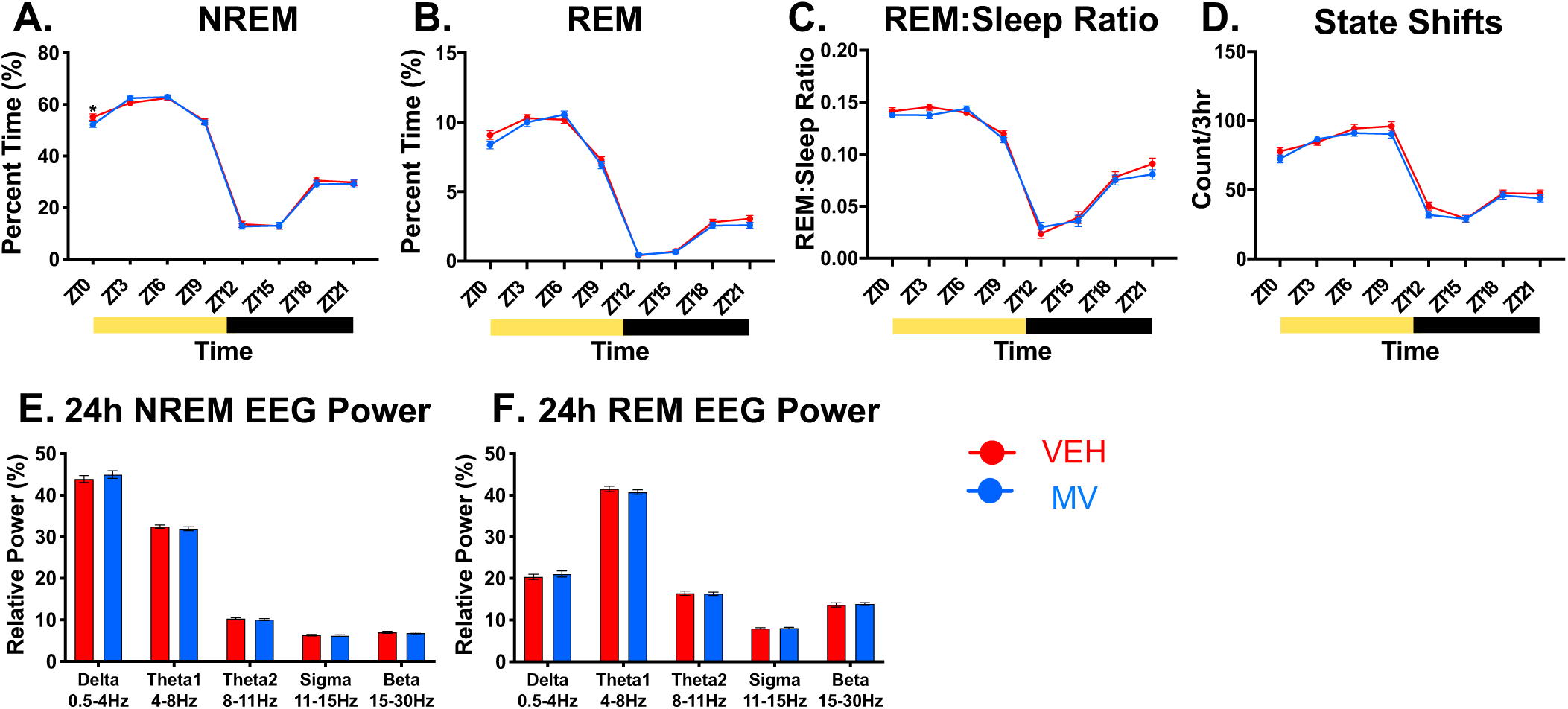
*M. vaccae* alone does not impact baseline sleep. Three days after the third injection of *M. vaccae* or vehicle, 24 h of baseline sleep was recorded. (A) NREM, (B) REM, (C) REM:Sleep, and (D) state shifts are reported in 3 hour bins. Yellow bars below the x axes represent times where the lights were on, while black bars represent times the lights were off. (E) 24 hour NREM and (F) 24 hour REM EEG power bands are expressed as a percent of total power. Data are mean + SEM. Symbols: \**p* < 0.05 (Tukey’s post hoc test). *n* = 52-53/group. Abbreviations: EEG, electroencephalography; MV, *Mycobacterium vaccae*; NCTC 11659 injection; NREM, non-rapid eye movement; REM, rapid eye movement; VEH, vehicle injection; ZT, zeitgeber time.

**Figure S2:**
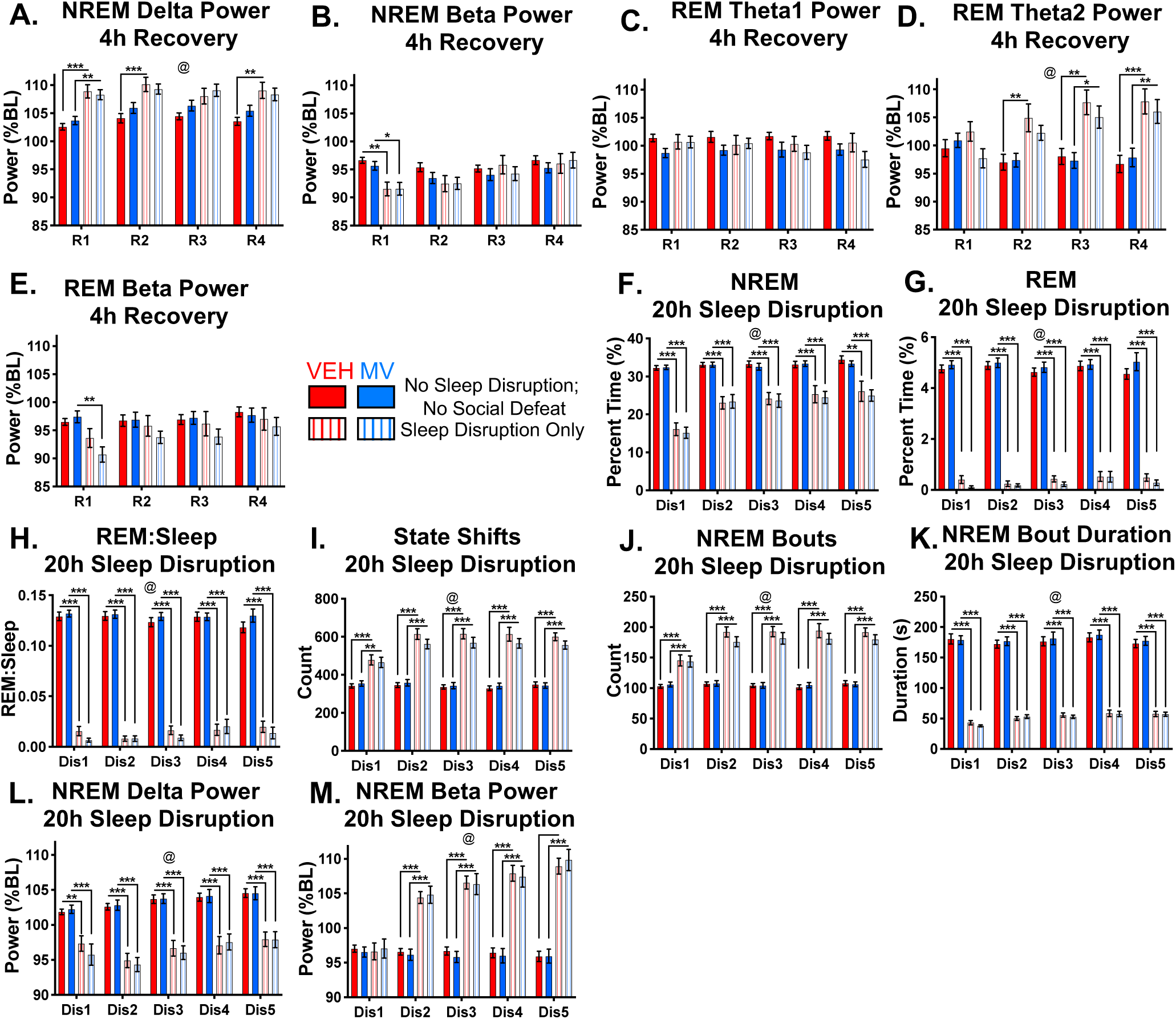
Other Sleep Measures During the Sleep Disruption Protocol. Continuation of Figure 1. (A) NREM EEG delta (0.5-4 Hz) power, (B) NREM EEG beta (15-30 Hz) power, (C) REM theta1 (4-8 Hz) power, (D) REM theta2 (8-12 Hz) power, and (E) REM beta power during the 4 hour (ZT2-ZT6) *ad libitum* sleep recovery windows during the sleep disruption protocol. EEG power is reported as a percent of baseline values between ZT2-ZT6. (F) NREM, (G) REM, (H) REM:Sleep ratio, (I) state shifts, (J) NREM bout count, (K) median NREM bout duration, (L) NREM EEG delta power, and (M) NREM EEG beta power during the daily 20-hour window (ZT6-ZT2) that the automated sleep disruption units were functioning. Data are means + SEM. Symbols: @ *p* < 1.0 x 10-4 (overall effect of “Sleep Disruption”), linear mixed effects model; \**p* < 0.05, \*\**p* < 0.01, \*\*\**p* < 0.001 (Tukey’s post hoc test). *n* = 29-30/group. Abbreviations: BL, baseline; Dis, sleep disruption; EEG, electroencephalography; MV, *Mycobacterium vaccae*; NCTC 11659 injection; NREM, non-rapid eye movement; REM, rapid eye movement; VEH, vehicle injection; ZT, zeitgeber time.

**Figure S3:**
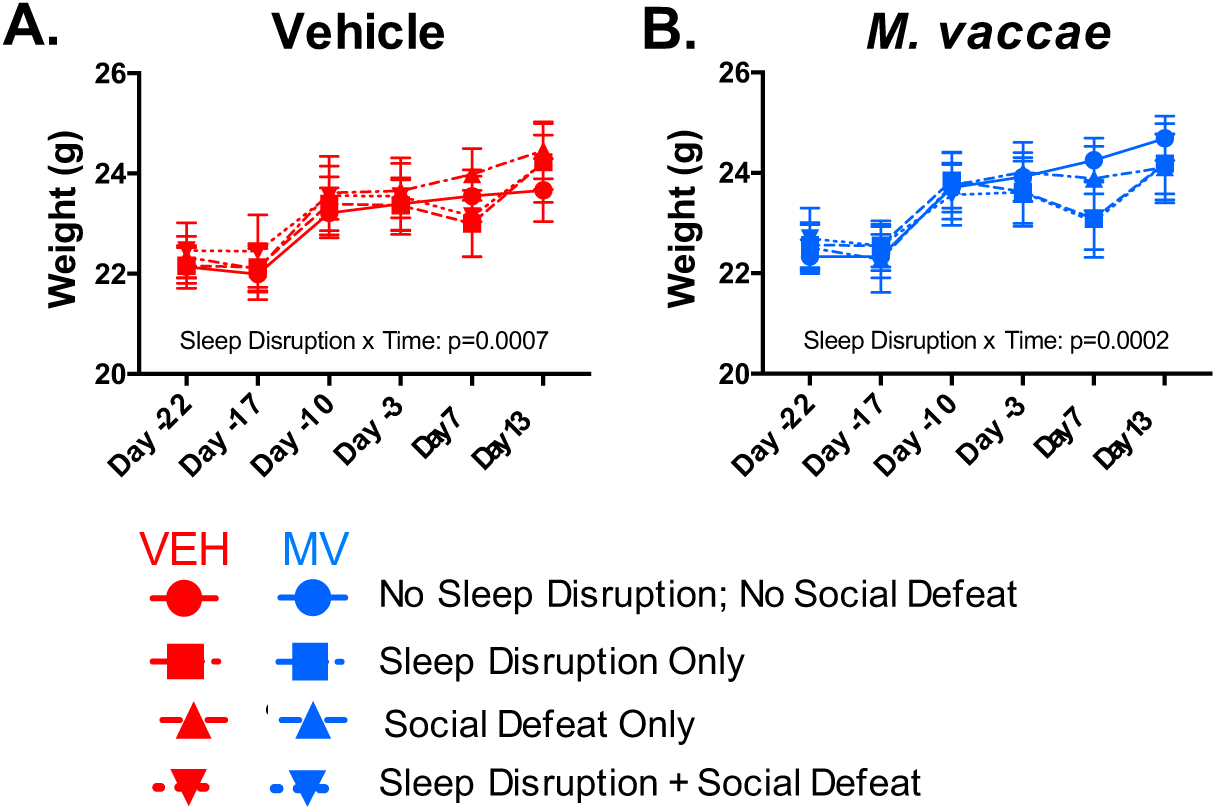
Body Weights Across the Experiment. Body weight data for all (A) vehicle and (B) *M. vaccae*-treated groups. Body weights were measured on the day of EEG/EMG surgery, on each injection day, on the day of OLM testing, and at the end of the experiment. Linear mixed effects model testing was performed for each treatment condition. Data are mean + SEM. *n =* 14-15/group. Abbreviations: EEG, electroencephalography; EMG, electromyography; MV, *Mycobacterium vaccae*; NCTC 11659 injection; OLM, object location memory; VEH, vehicle injection.

**Figure S4:**
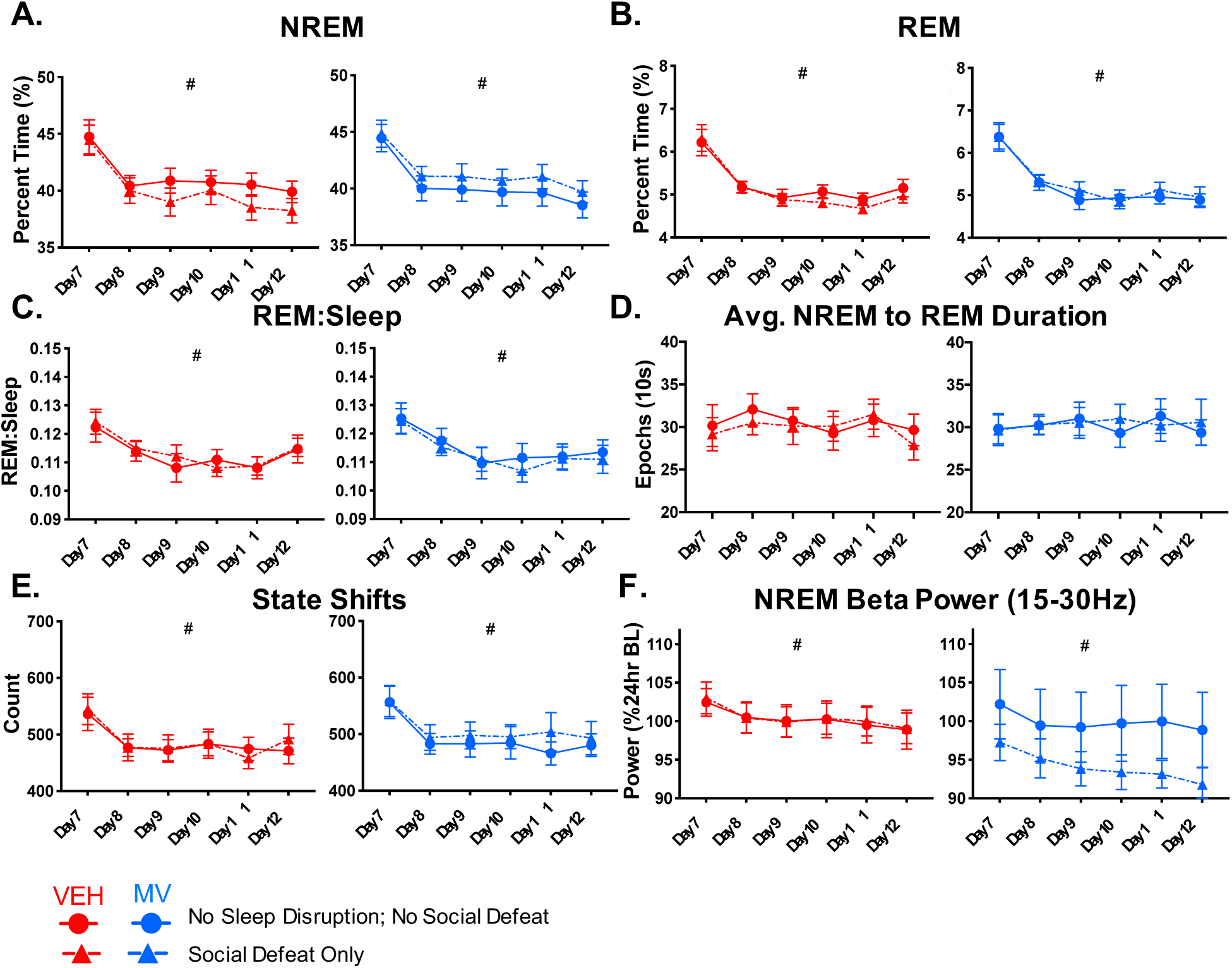
Social Defeat Alone Did Not Result in Lasting Sleep Changes. Continuation of Figure 3. (A) NREM, (B) REM, (C) REM:Sleep ratio, (D) average number of epochs of NREM preceding a REM bout, (E) state shifts, and NREM EEG beta (15-30 Hz) power is reported for control and social defeat alone groups. Symbols: # *p* < 0.05 (overall effect of time), linear mixed effects model. Data are mean + SEM. *n* = 11-14/group. Abbreviations: MV, *Mycobacterium vaccae*; NCTC 11659 injection; NREM, non-rapid eye movement; REM, rapid eye movement; VEH, vehicle injection.

**Figure S5:**
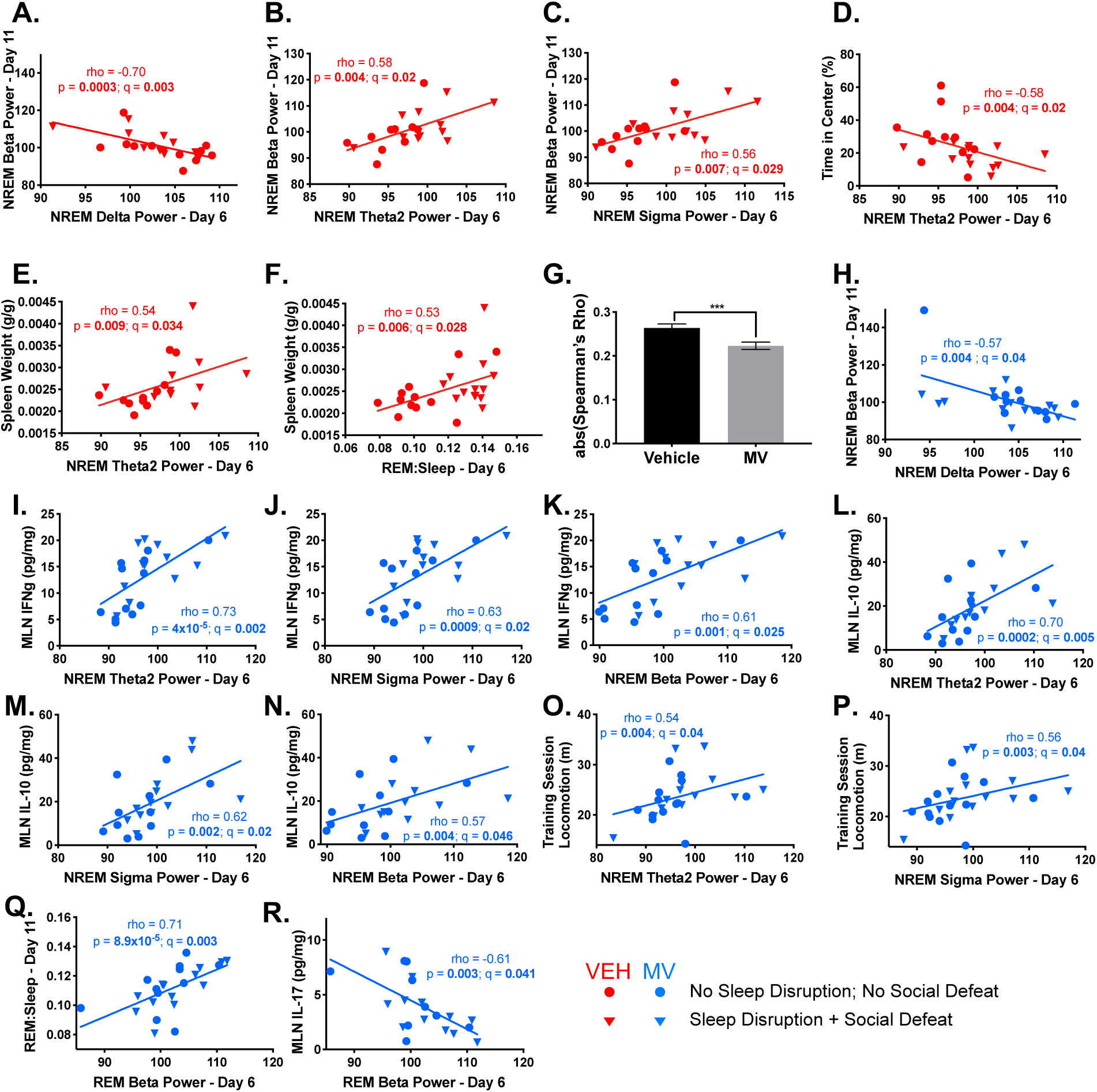
Selected Associations Between Acute Sleep Changes, Lasting Sleep Changes, Behavior, and Physiological Measures. Continuation of Figure 6. Pairwise Spearman’s rank order correlations of 32 measures of interest from throughout the experiment were computed for vehicle-treated and *M. vaccae* (MV)-treated mice that received either control manipulations or sleep disruption plus social defeat (double hit). NREM beta (15-30 Hz) EEG power on Day 11 (percent of baseline) vs (A) NREM delta (0.5-4 Hz) power (B) NREM theta2 (8-12 Hz) power, and (C) NREM sigma (12-15 Hz) power on Day 6 in vehicle-treated mice. (D) Time spent in the center of the arena during the object location memory (OLM) training session vs NREM theta2 power on Day 6 in vehicle-treated mice. Spleen weight at the end of the experiment vs (E) NREM theta 2 power on Day 6 and (F) REM:Sleep ratio on Day 6 in vehicle-treated mice. (G) The average of the absolute value of all pairwise Spearman’s Rho (*n =* 512) values for the vehicle and MV treatment groups. (H) NREM beta power on Day 11 vs NREM delta power on Day 6 in MV-treated mice. Mesenteric lymph node (MLN) interferon gamma (IFNγ) vs (I) NREM theta2 power on Day 6, (J) NREM sigma power on Day 6, and (K) NREM beta power on Day 6 in MV-treated mice. MLN interleukin (IL) 10 vs (L) NREM theta2 power on Day 6, (M) NREM sigma power on Day 6, and (N) NREM beta power on Day 6 in MV-treated mice. Locomotion during the training session of the OLM task vs (O) NREM theta2 power on Day 6, and (P) NREM sigma power on Day 6 in MV-treated mice. (Q) REM:Sleep ratio on Day 11 vs REM beta power on Day 6 in MV-treated mice. (R) MLN IL-17 vs REM beta power on Day 6 in MV-treated mice. Symbols: \*\*\**p* < 0.001, Wilcoxon Rank-Sum test. *n* = 9-12/group. Abbreviations: IL, interleukin; MLN, mesenteric lymph node; MV, *Mycobacterium vaccae*; NCTC 11659 injection; NREM, non-rapid eye movement; OLM, object location memory; REM, rapid eye movement; VEH, vehicle injection.

**Table S1: Complete Statistical Results for Figures 1-5, S1-S4**. Spreadsheet containing results of all linear mixed effects modeling/ANOVA performed in the experiment, along with post hoc comparisons and *p* values. Each tab of the spreadsheet contains the results for a different figure.

**Table S2: Results of correlation studies depicted in Figures 6 and S5**. Spreadsheet containing the Spearman’s rho, *p* value, and *q* value matrices resulting from the Spearman’s Rank Based correlation analysis performed between sleep, behavior, and physiological data.

